# Short Linear Motifs In Intrinsically Disordered Regions Modulate HOG Signaling Capacity

**DOI:** 10.1101/199653

**Authors:** Bob Strome, Ian Hsu, Mitchell Li Cheong Man, Taraneh Zarin, Alex Nguyen Ba, Alan M Moses

## Abstract

The effort to characterize intrinsically disordered regions of signaling proteins is rapidly expanding. An important class of disordered interaction modules are ubiquitous and functionally diverse elements known as short linear motifs (SLiMs). To further examine the role of SLiMs in signal transduction, we used a previously devised bioinformatics method to predict evolutionarily conserved SLiMs within a well-characterized pathway in S. cerevisiae. Using a single cell, reporter-based flow cytometry assay in conjunction with a fluorescent reporter driven by a pathway-specific promoter, we quantitatively assessed pathway output via systematic deletions of individual motifs. We found that, when deleted, 34% (10/29) of predicted SLiMs displayed a significant decrease in pathway output, providing evidence that these motifs play a role in signal transduction. In addition, we show that perturbations of parameters in a previously published stochastic model of HOG signaling could reproduce the quantitative effects of 4 out of 7 mutations in previously unknown SLiMs. Our study suggests that, even in well-characterized pathways, large numbers of functional elements remain undiscovered, and that challenges remain for application of systems biology models to interpret the effects of mutations in signalling pathways.

**One-sentence Summary:** Mutations of short conserved elements in disordered regions have quantitative effects on a model signaling pathway.

## Introduction

There is an increasing proportion of known signaling interactions mediated by regions of proteins lacking a defined tertiary structure under native conditions, known as intrinsically disordered regions (IDRs) (*1*, *2*). Comparative proteomics studies have used evolutionary conservation to identify short functional elements in these intrinsically disordered regions,(*3*) and suggest that many remain undiscovered (*4*). The short conserved elements identified are often short linear motifs (SLiMs), which are ubiquitous and functionally diverse elements (3-10 amino acids), and are often (∼80%) located within the IDRs (*5*). Unlike well-defined large, globular domains, the functional significance of these motifs is inherently difficult to determine due to their small size and relatively low binding affinities (*6*).

One of the best-understood functions of SLiMs is to mediate signaling interactions, both through post-translational modifications and peptide-domain interactions (*1*, *2*, *7*–*9*). Recent evidence also suggests "weak" SLiMs in disordered regions can facilitate/recruit post-translational modifications (*10*). Components of well-characterized model signaling pathways, such as the high osmolarity glycerol (HOG) MAPK pathway in budding yeast (*11*–*15*), contain numerous short conserved elements (predicted short linear motifs, pSLiMs) within their IDRs. This suggests that, even in well-studied pathways, there are still many uncharacterized elements to be discovered. Here we set out to test this prediction.

Because each SLiM is thought to mediate a single, usually transient interaction, we expect mutations in them to have quantitative effects on protein function. We therefore decided to employ a quantitative, single cell reporter assay (*16*, *17*) to detect subtle differences in activation of the pathway. Since the HOG pathway has been the subject of detailed modeling (*18*–*22*), we attempted to interpret the effects of mutations within mechanistic, quantitative models.

Overall, we identified 7 previously uncharacterized short protein sequences (between 3 and 17 amino acids in length) that significantly impact pathway output when deleted. Among these and three previously known motifs, we observed a wide quantitative range of effects on pathway output and tested whether these effects could be explained by perturbations of parameters in a previously proposed quantitative model of the HOG pathway (*21*). We found that the model can quantitatively reproduce the effects of 5 of 10 of the mutations, but in general it cannot identify the part of the pathway affected by the mutation. For mutations with the largest effects, the model agrees with the data only qualitatively. Although it is increasingly appreciated that disordered regions mediate important signaling interactions, our study highlights that systematic approaches are needed to identify the large number and quantitative effects of these interactions. We suggest that interpretation of the quantitative effects of mutations using mechanistic models is an important area for further research in systems biology.

## Results

### Well-characterized signaling pathways contain short conserved elements in disordered regions

We searched for short motifs in disordered regions in model signaling pathways in yeast using previously described methods (*4*). The HOG pathway is a well-characterized and highly conserved MAPK pathway (*11*–*15*). In budding yeast this pathway responds to increased external osmolarity, altering metabolism, cell cycle progression and gene expression to re-establish cell homeostasis. The pathway consists of three MAPKK’s comprising two partially redundant incoming branches (Sln1-dependant and Sho1-dependant branches)(*23*) that function in parallel, eventually converging at the MAPKK (Pbs2)(*12*–*14*, *18*). Activated Pbs2 in turn phospho-activates Hog1p which then translocates to the nucleus, initiating transcription of up to 600 genes (*12*, *13*). Here we focus on the Sho1-dependant branch of the pathway which can be studied in isolation by deleting the two redundant MAPKK’s of the Sln1-dependant branch (*24*, *25*).

Analysis of the Sho1-dependant branch of the HOG pathway yielded 29 predicted short linear motifs (pSLiMs) within 14 disordered regions (encompassing 147 residues, or 1.9% of the amino acids in the proteins in this branch of the pathway, Figure 1). Among these short conserved elements, at least 4 correspond to known short linear motifs in this pathway. Because the HOG pathway (particularly the Sho1 branch) has been the subject of extensive molecular characterization (*26*, *27*) and detailed quantitative analysis and modeling (*12*, *13*, *18*), we were surprised to identify 24 short conserved elements in disordered regions for which we could not identify functions in the literature. To confirm that the HOG pathway was not unusual in this regard, we examined another well-characterized yeast pathway, as well as a human pathway, and observed similar conserved putative motifs in both. The cell wall integrity pathway in yeast (*23*) contains 36 total pSLiMs in 11 proteins (Table S1A), and the human canonical Wnt regulatory pathway (*28*) contains 23 pSLiMs in 14 proteins (pSLiMs predicted in human (*29*), Table S1B). This indicates that many signaling pathways may contain large numbers of uncharacterized short linear motifs (see discussion).

**Figure 1:**
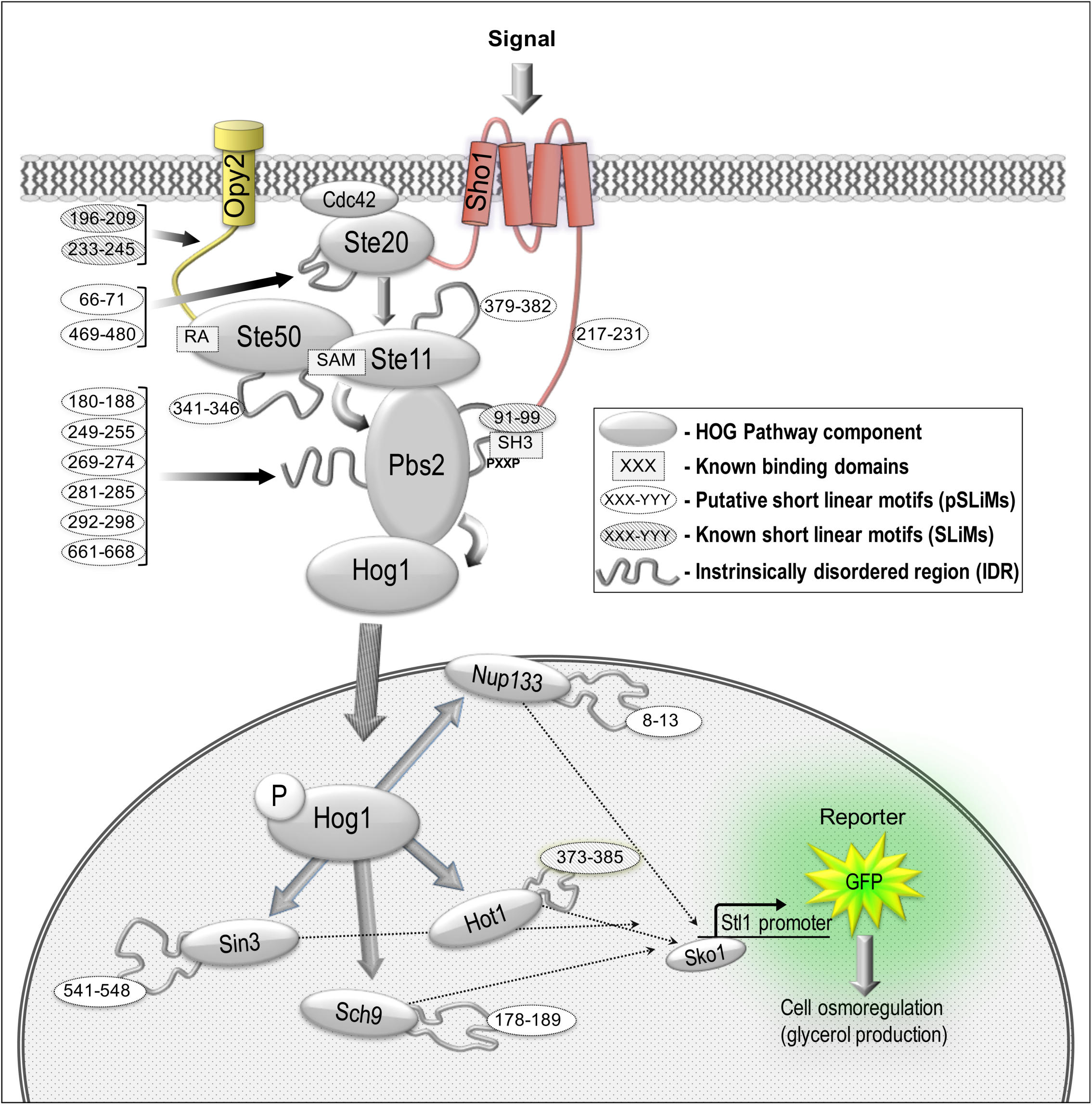
Schematic diagram of the Sho1-dependant branch of the yeast high osmolarity glycerol (HOG) pathway as a model system for analysis of short linear motif function in signal transduction. Upon exposure to high osmolarity, the SH3 domain of the membrane-bound osmosensor Sho1p binds to PXXP motif in the MAPK kinase (MAPKK) Pbs2p, initiating a phosphorylation cascade (grey arrows) that culminates in phosphorylation of the MAPK Hog1p. Activated Hog1p-P then accumulates inside the nucleus and initiates a complex transcriptional response in order to re-establish homeostasis within the cell. A Hog1-responsive promoter driving yeGFP reporter expression was used to quantify pathway activity for a series of deletions in predicted short linear motifs in pathway components – deleted residues shown in ovals for each module.

### Mutations in known short motifs affect HOG signaling

We sought to test for functional importance of the short conserved elements (pSLiMs) in the HOG signaling pathway. We first confirmed that previously characterized pSLiMs in the HOG pathway affected signaling function in a reporter-based flow-cytometry assay that can quantitatively measure pathway activity using a fluorescent reporter downstream of Hog1-specific transcriptional output (see Methods). We made deletions and point mutations in the conserved Sho1 SH3 domain-binding site in Pbs2 (the MAP2K, Figure 1) (*26*, *27*), as well as deletions in a known motif in Hot1 (a downstream transcription factor)(*30*), a known Ste50 binding site in Opy2 (membrane anchor for Ste50p) (*31*) and the Bem1 binding site in Ste20 (*32*).

We found that three of these four known motifs showed reduction in reporter activity (Figure 2). This confirms that our assay has power to detect the effects of mutations of known signaling interactions mediated by short protein sequences in disordered regions. Deletion of the proline-rich SH3-binding site in Pbs2 (*26*) virtually abolished the signaling response (Figure 2a). To confirm that the strong effect observed in the Pbs2 deletion was due to the specific signaling interaction, and not a global effect on protein function due to deletion of the binding site, we also measured the signaling response in the presence of two point mutations (residues P96 and P99) that were previously shown to compromise binding (*26*). As expected, signaling in this two-point proline mutant showed a signaling response that was similar to that of the binding site deletion (Figure 2A).

**Figure 2:**
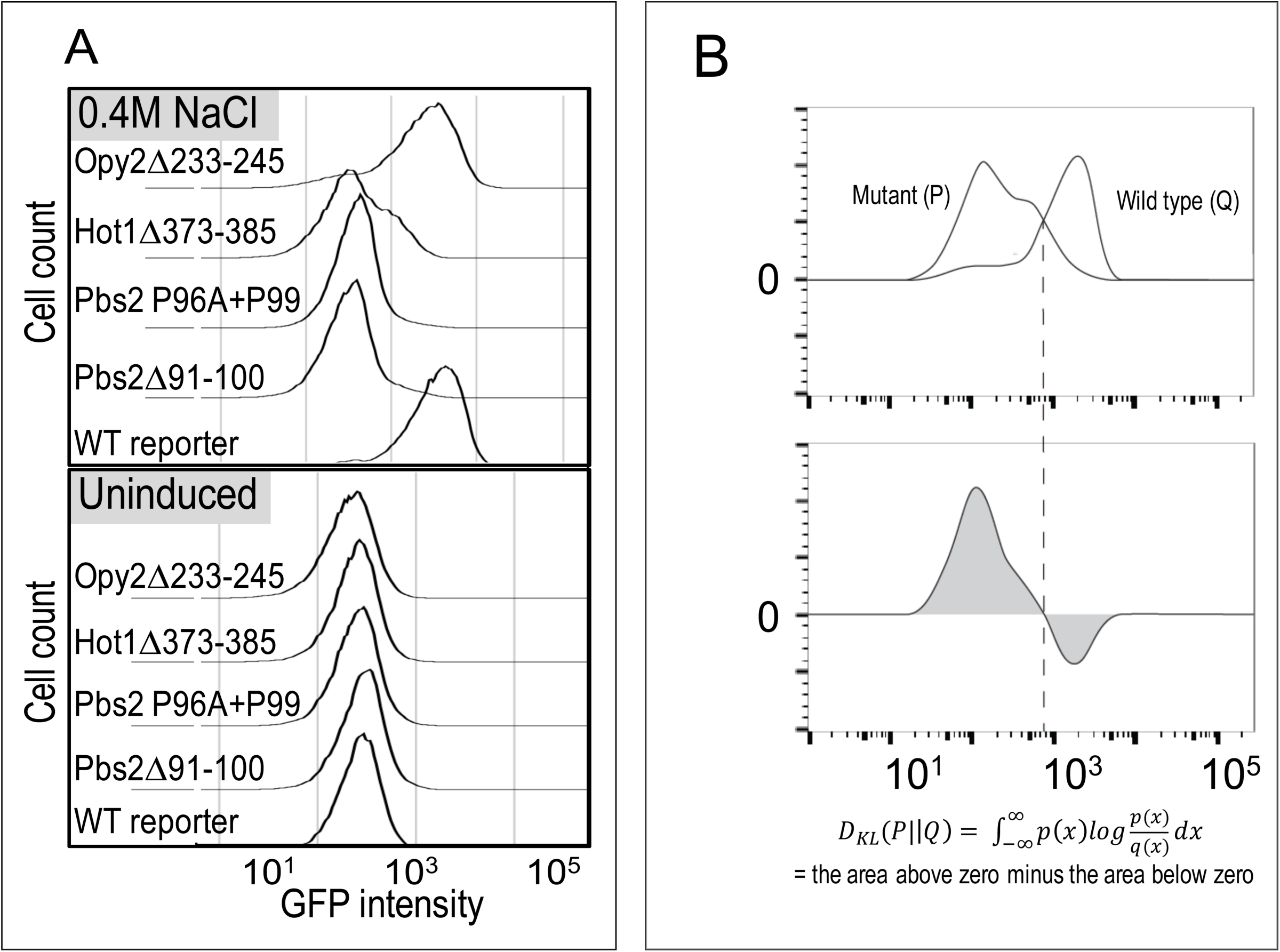
Deletions of known short linear motifs impairs HOG pathway-specific reporter activation. **A.** Histograms of flow cytometry data showing expression of GFP driven by a Hog1-specific promoter (Stl1) before (lower panel) and after (upper panel) a 60 minute induction with 0.4M NaCl. Pbs2: both a 10 residue deletion (Δ91-100) and double point mutant (P96A + P99A) of SH3 binding domain essentially eliminates reporter output; Hot1: (glycerol biosynthesis transcription factor) 13 residue deletion of a known activation domain (373-385) substantially impairs pathway output and effects a bimodal distribution; Opy2: 13 residue deletion of known Ste50 binding domain (233-245) effects a partial reduction in reporter output. **B.** Comparison of distributions using the KL divergence, *D_KL_*(*P||Q*). The top panel shows cartoon distributions representing either mutant (P) or wild-type (Q) flow cytometry data. The bottom panel shows the log ratio of the two distributions weighted by the probability under the mutant. The KL divergence (equation below the bottom panel) is the integral of this curve (grey area). Dashed line shows that when P(x) and Q(x) have the same value, there is no contribution to KL divergence.

Because our flow cytometry assay measures reporter activity in single cells, we compared the distribution of reporter activity measurements for mutant strains to wild-type using the Kullback-Leibler Divergence (KL divergence, see methods). If two distributions are identical, the measure gives a value of zero; any real data will have positive KL divergence because of experimental (and possibly biological) noise. To understand how much KL divergence we could expect due to these (non-genetic) sources, we compared the distribution of wt reporter measurements to themselves (see methods), and found on average KL= 0.016 (+/- 0.014 standard deviation). To assess whether a mutant affected HOG signaling, we compared the divergence (in replicate experiments) of the mutant distribution to wt. If the divergence of the mutant to the wt was significantly greater than the wt compared with itself, we consider that mutant to impact signaling.

Using this approach, we found the deletion of a known Ste50 binding site in Opy2 showed a small, but statistically significant quantitative effect (*P*<0.001, Table S2, Figure 2) on reporter expression. This is notable, as this binding site is one of three cooperative motifs that are required for the full Ste50-Opy2 interaction (*31*). Indicating that our quantitative assay may be sensitive enough to identify individual components of other cooperative binding interactions.

Unlike the binding site in Opy2, deletion of the known Bem1 binding site in Ste20 (*32*) showed no significant effect on reporter expression (Table S2). This motif is known to be involved in yeast mating signaling (*32*), and its functional effect would likely be detected using a different assay. This shows that our assay cannot be expected to detect all functional motifs in HOG pathway components (see discussion).

Perturbation of Hot1 activity is known to lead to stochastic responses in the HOG pathway (*33*). Interestingly, and consistent with previous studies implicating Hot1 with stochastic responses, we found that deletion of the known Hog1 binding region in Hot1 (residues 373-385; Alepuz et al., 2003) led to a bimodal response (P<0.00001, Table S2, Figure 2), where some of the cells showed no difference from the uninduced state, while others showed a weak response (Figure 2). These results confirm that our assay can detect both quantitative, population level as well as stochastic, single cell level effects on signaling. Furthermore, these results suggest that mutation of a short conserved binding site can enhance stochastic responses (see discussion).

### 7/14 previously unknown short conserved elements in disordered regions quantitatively decrease HOG signaling

Using the same assay as above, we tested a cohort of previously uncharacterized pSLiMs in the disordered regions of the HOG pathway (Figure 1). Of the predicted novel motifs within the pathway, we selected 15 candidates of at least 4 residues for functional analysis based our examination of the alignments with preference given to pSLiMs in the kinases and other signaling proteins. We created 14 yeast strains in which we deleted an individual pSLiM and quantified pathway output (Table S2). For one of the mutants (Sch9 Δ279-95) we were not able to obtain strains with consistent pathway output, apparently due to secondary mutations. We therefore excluded it from further analysis. Of the remaining 14 strains, 7 (50%) showed changes in the reporter distribution (P-value<0.05 after Bonferroni correction; Figure 3).

**Figure 3:**
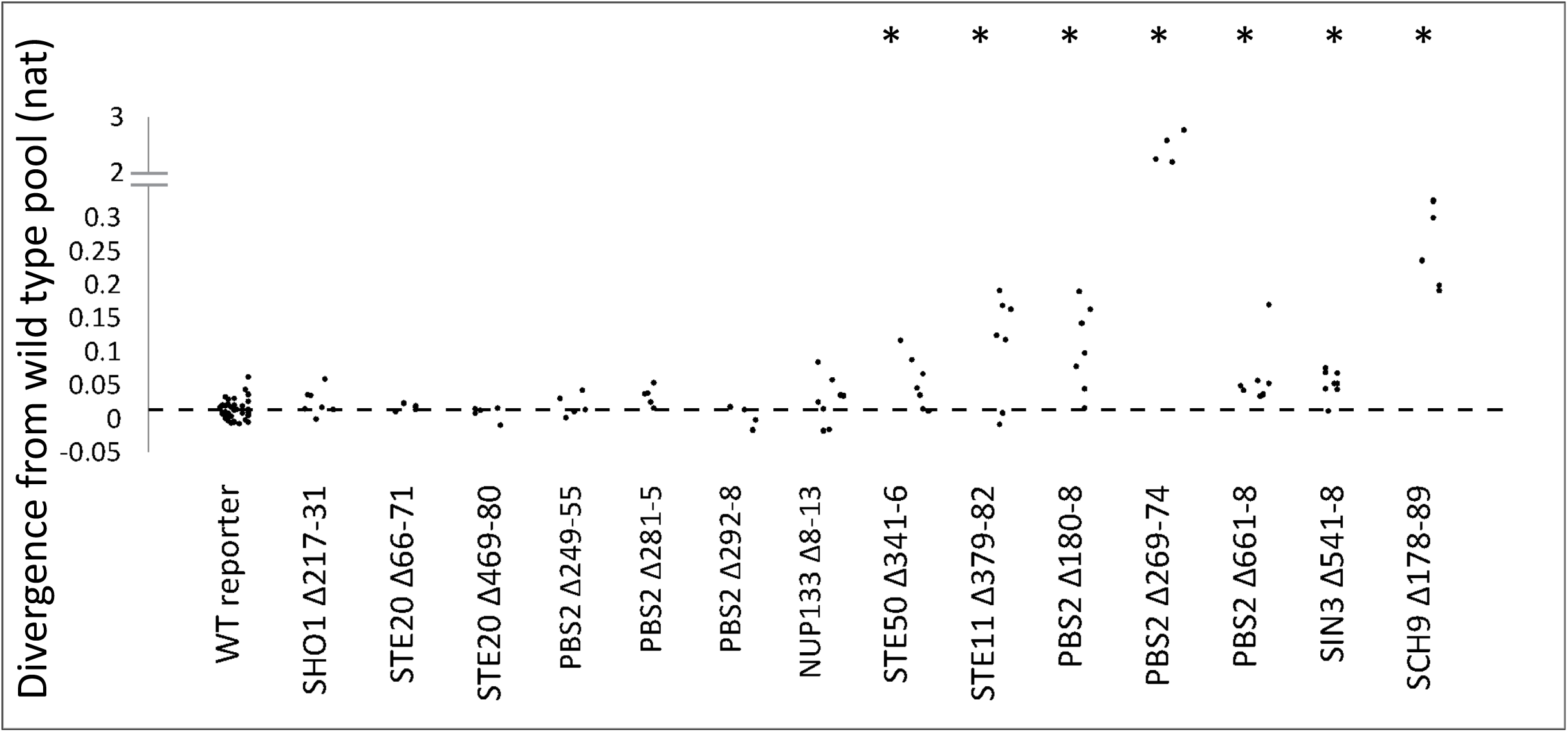
7/13 previously unknown short conserved elements in disordered regions show significantly different reporter expression distributions compared with wild type. Each point shows the KL divergence of a single replicate from the wild type pool. pSLiM deletion strains are indicated along the horizontal axis. Dashed line indicates the average divergence of the wt replicates from the wt pooled data (see text for details). Asterisks indicate the strains that have significantly larger KL divergences than wild type reporter (Wilcoxon Rank Sum Test P<0.05).

These varied effects include nearly complete abolishment of the pathway activity for deletion of a previously uncharacterized proline-rich sequence in Pbs2, while most mutants showed subtle quantitative effects. The large effect of the mutation in the Pbs2 pSLIM is consistent with Pbs2’s role in localizing the signaling complex (*13*, *27*, *31*), but we were surprised that this seemingly critical motif was not identified before. That most of the pSLIMs show quantitative effects is consistent with our expectation that they are involved in transient, possibly combinatorial regulatory interactions, much like the known motifs in Opy2. Taken together, this data indicates that the novel motifs show a varied functional significance similar to the previously known motifs (Figure 2).

### Comparison of SLiM mutants to a stochastic model of HOG signaling

In principle, quantitative models of signaling can be used to connect genotype to phenotype, and could therefore be used to predict the impact of mutations in disordered regions. Considering that signaling proteins are often mutated in human disease, this represents a potentially important area of application for quantitative models.

The yeast MAPK signaling pathways have served as a foundation for the understanding of signaling in more complex organisms (*23*), and the HOG pathway in particular has been the subject of several mechanistic modeling studies (*18*, *20*–*22*, *34*–*36*). Previous modeling studies of the HOG pathway have demonstrated that the distribution of quantitative single cell reporter measurements can be explained using stochastic simulations of models of signaling and gene expression (*20*, *21*, *36*). However, few studies have explored the extent to which these models generalize to new datasets, nor have they considered whether models can capture quantitative effects of mutations.

We therefore first tested whether a previously published model of HOG signaling (*21*) could explain the reporter expression we observed for the wildtype strain. We simulated individual cell trajectories and normalized the output of the model to convert the scale (in molecules) to fluorescence and computed the KL divergence between the distribution predicted by the model and the real data. We found that the model at stress level 18 (with other parameters unchanged) could quantitatively reproduce the wild-type data observed in reporter assay (Figure 4a), such that the KL divergence between the model and the observed data was 0.061 (+/- 0.03431 standard deviation), which is larger, but on the same order as the divergence between replicates (0.016 +/- 0.014). This indicates that the previously developed model (with appropriate normalization) can explain the distribution of reporter expression in our experimental setup.

**Figure 4:**
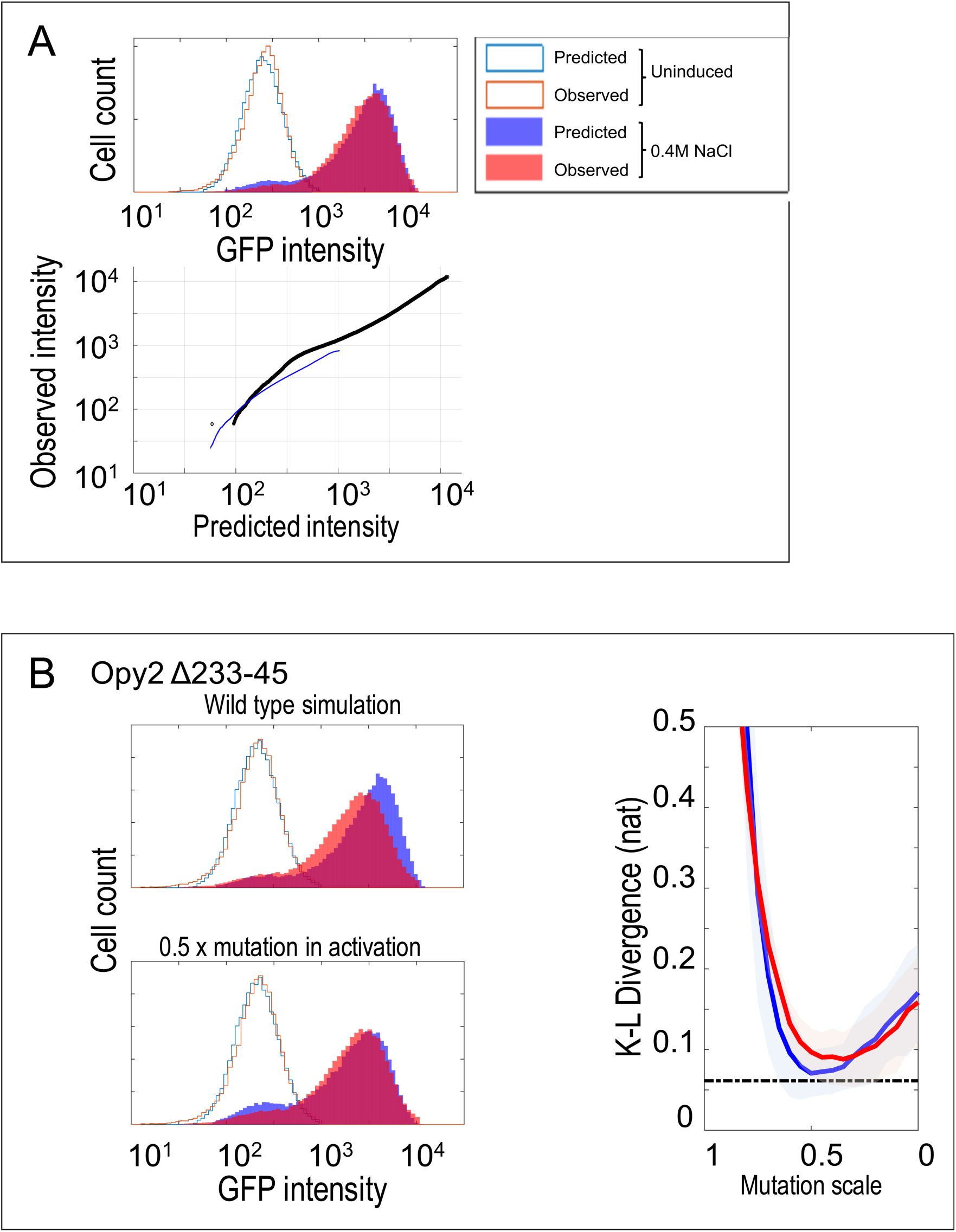
Simulations of a stochastic model can generate histogram data that resemble wild type and mutant distributions. **A.** A representative comparison between data from a wild type experimental replicate (red traces) and the prediction of the model (blue traces). The lower panel shows a QQ plot summarizing the similarity of the two distributions by comparing each quantile. The blue line indicates comparison of the uninduced distributions (unfilled histograms in top panel), while the black line indicates comparison of the distributions after stimulation for 60 minutes in 0.4 M NaCl (filled histograms in the top panel) **B.** Comparison of an experimental replicate of Opy2Δ233-45 (red traces) with predictions of the model (blue traces). The top left panel shows comparison with the wild type model, while the bottom left plot shows the same mutant data compared to the prediction of the simulation when the value of the activity parameter is decreased by 50% (indicated as 0.5 x). The plot on the right summarizes how K-L divergence changes while parameter values decrease. Red lines and blue lines respectively represent the K-L divergence as the remodelling parameter and activity parameter are decreased. The shaded areas show +/- 3 times the standard errors. The dashed line is mean of wild type divergence from the wild type simulation. Compared to the histogram of induced mutant, the predicted histogram simulated with original parameters shifts toward higher intensity.

To predict the effect of mutations in the pathway, we assumed that each mutation would affect only one of the 5 parameters associated with signaling components in the model (i.e., we did not consider parameters that control global rates of transcription or translation because our mutations were made within the signaling components). Because the simplified model does not explicitly include signaling components (or their protein interactions) we do not know in advance which parameter each mutation would affect, nor do we know the magnitude of the effect the mutation in a SLiM would have on the parameter. However, through simulations of where the different parameters were reduced individually (to model the effects of mutations), we noticed that the effects on the parameters fell into two groups (Supplementary figure 1), based on whether the parameters affected the pathway activation steps before Hog1 (which we refer to as activity parameters) or the slow transcriptional steps below Hog1 (which we refer to as remodelling parameters). This apparent redundancy in the effects of perturbations to the parameters greatly simplified our analysis of mutant pathways: we need only compare our data to two sets of perturbed simulations. Nevertheless, for each type of parameter, we don’t know what quantitative effect the pSLIM mutation will have.

To evaluate the fit of the perturbed model to the reporter expression data from the mutant signaling pathways, we identified the parameter value that minimized the KL divergence between the mutant data and the prediction of the perturbed model. For example, for the known binding site in Opy2, we found that a model where the activity parameter was reduced by 50% fit the mutant data with KL divergences comparable to the fit of the unperturbed model to the wt data (Figure 4b). Indeed, the KL divergences between this perturbed model and the Opy2 mutant data were significantly smaller than those between the Opy2 mutant data and the wt model (P=0.02, Bonferroni corrected Rank Sum test) which is expected in part because the mutant model has added flexibility. Interestingly, we could not achieve a significantly better fit for the models when the remodelling parameter was perturbed (P=0.4, Bonferroni corrected Rank Sum Test) suggesting that the mutation in the Opy2 binding site affects the signal activation upstream of Hog1, which is consistent with the known position of Opy2 in the pathway (Figure 1). To test this hypothesis directly, we compared the divergences of the best perturbed activity parameter model to the best perturbed remodelling parameter model, but we found that the divergences between the data and the two perturbed models were not significantly different. Nevertheless, this analysis suggests that perturbations of single parameters in the HOG model can explain the quantitative effects of the pSLiM deletion mutants.

We applied this analysis systematically to test which other mutant data could be explained by the model. For four of the seven previously unknown pSLiM deletion mutants that had significant effects on pathway output (shown in Fig.3), we found the fit of the perturbed model to the mutant data was comparable to the fit of the wild type model to the wild type data (Figure 5). Considering that this model was not designed to fit quantitative data, nor to model effects of mutations (*21*), this generalization capacity is impressive. For the remaining three mutant strains the divergences of the perturbed model were significantly larger. We noted that these three strains represented the mutants with the largest effects (see discussion). We attempted to use the fits of the model to predict which parameter the pSLiM mutation affected, and although in some cases the fit of a perturbed model with one parameter or the other was slightly better, these differences were not significantly different (eg. Fig.6A, See discussion).

**Figure 5:**
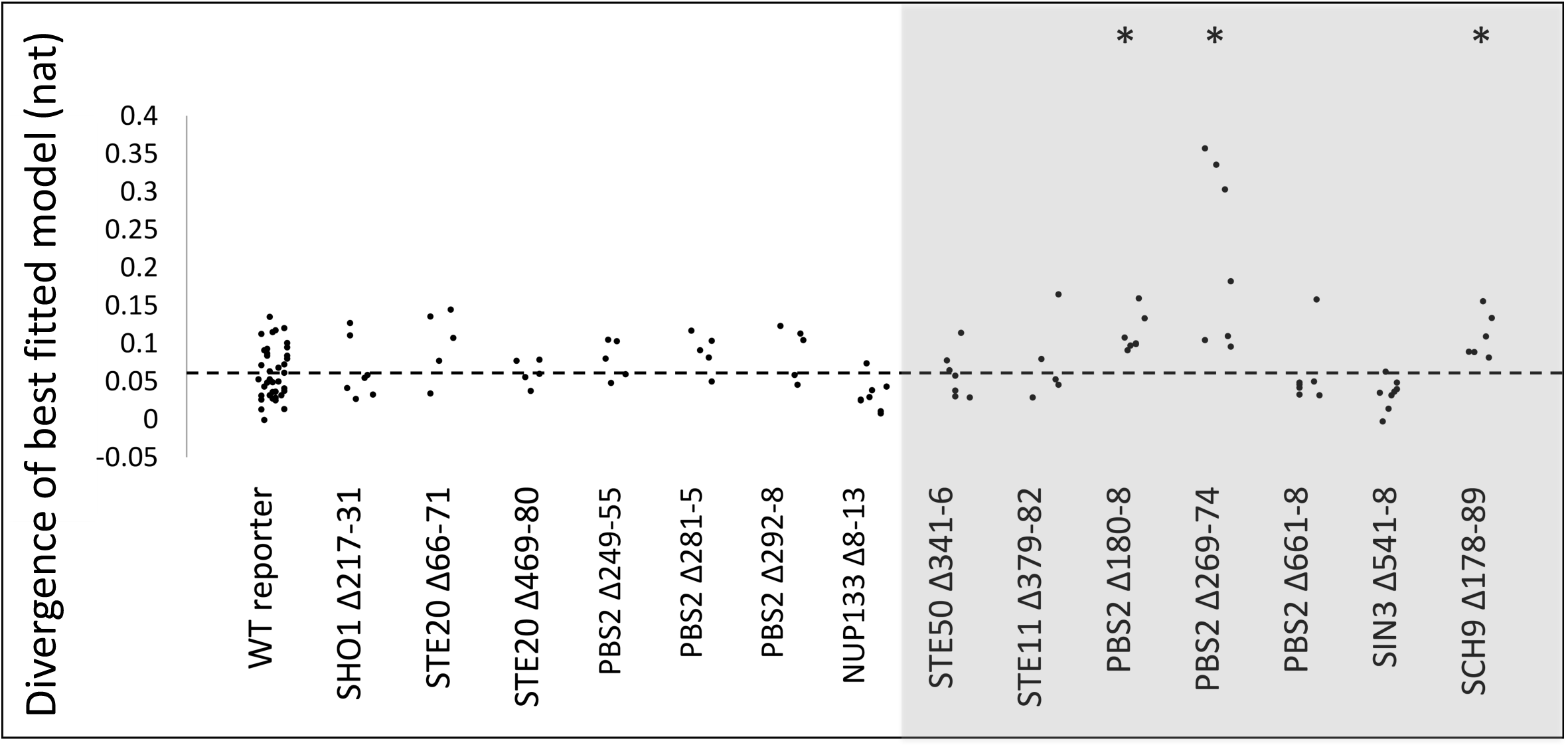
Comparisons of KL divergence of best fitting models. Each point represents the fit of one replicate to the best fitting model for that strain. Grey bar indicates strains whose reporter expression distribution was significantly different from wild type (Fig.3). Stars indicate strains whose KL divergence to the best fitting model is significantly worse than the wild type fit (WT reporter). Dashed line represents the mean of KL divergence of wild type replicates to the wild type model.

**Figure 6:**
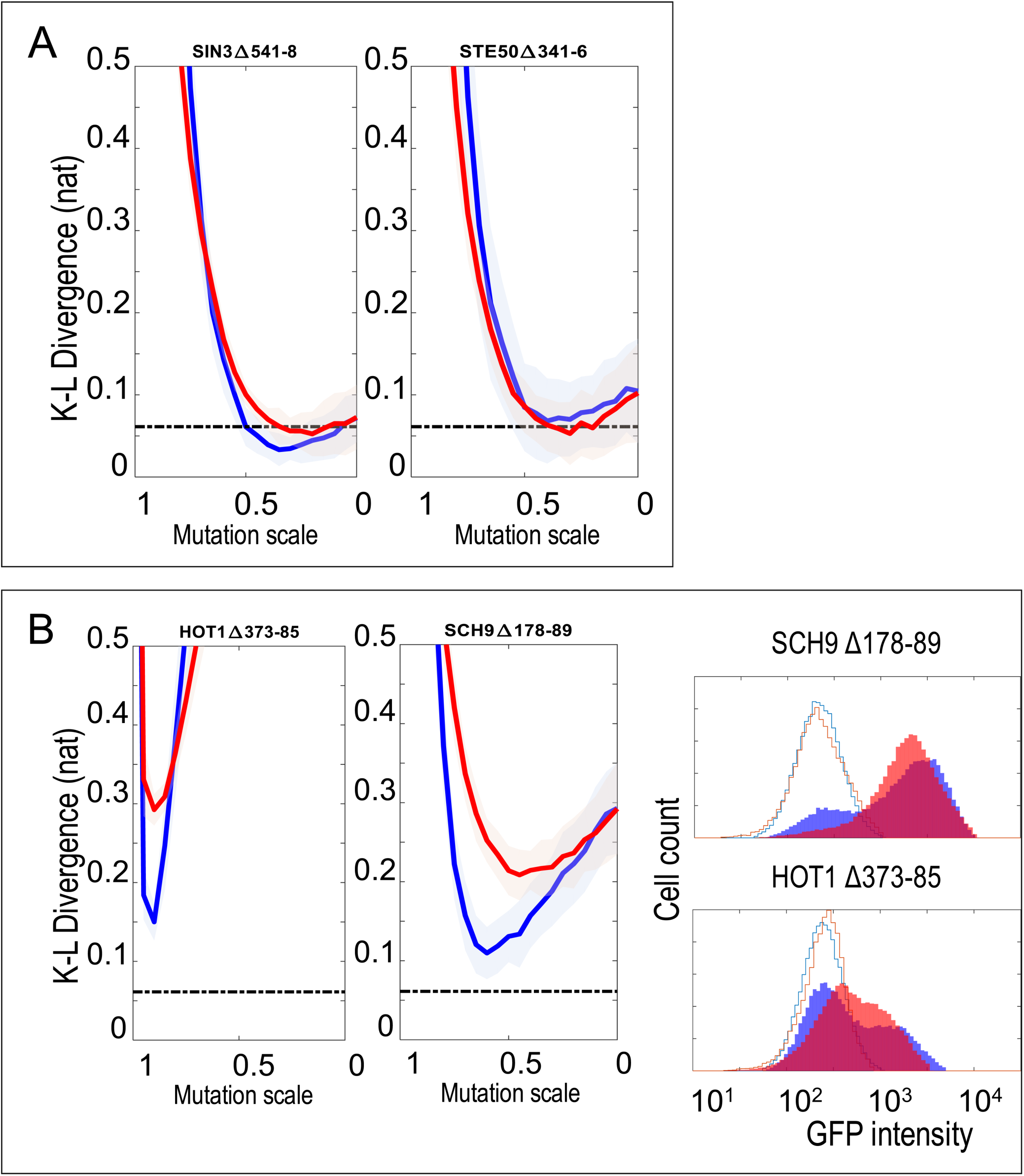
Fitting the stochastic model to wild type and previously unknown signaling mutants. **A.** examples of mutants that the model can explain. Red lines and blue lines respectively represent the K-L divergence as the remodelling parameter and activity parameter are decreased. The shaded areas show +/- 3 times the standard errors. The dashed line is mean of wild type divergence from the wild type simulation. **B.** Examples of mutants that the model cannot explain. The red and blue lines do not come close to the dashed line. The right panel shows comparisons of experimental replicates (red traces) to predictions of the model (blue traces) when the value of the activity parameter is decreased chosen to maximize the fit (minimize KL divergence).

We also noted for mutations with large effects, the model tended to over predict the population variation. For example, for Sch9 and Hot1 mutants, the best fitting models produce bimodal distributions where the two peaks are more separated than observed (Figure 6B). Further, for these mutations, the parameter that gives the best fitting mutant model is the activity parameter, which is inconsistent with the known roles of these proteins in the transcriptional part of the pathway.

Thus, by perturbing the parameters in the model to fit the pSLIM mutant data we can identify changes in parameters that improve the fit of the model to the data. However, it does not imply that the model is providing mechanistic insight into to the impact of the mutation on signaling function (see Discussion).

## Discussion

The use of stochastic models in systems biology is still a largely qualitative endeavour, consistent with recent research efforts indicating that fitting stochastic models to data remains a challenge in systems biology (*20*, *37*). To fit a previously published model (*21*) to our data, we assumed that mutations quantitatively effect a single parameter, performed simulations assuming all possible values of that parameter, and chose the value that minimizes the KL divergence of the simulation from data. Simulations revealed that many of the parameters had similar effects on the output pathway, suggesting that these parameters cannot be inferred from our experimental data (*38*). Encouragingly, we found that perturbed versions of the previously reported model could explain the distribution of signaling output. However, in general, we could not infer which parameter was affected by a given mutation, as both parameters were usually able to maximize the fit of the model to the data. This suggests that interpretation of mechanisms of mutations using systems biology models is an area requiring further research.

In addition, we noted that the model was better able to match the distributions for mutations that had small effects, and even for these mutants, the perturbation to the parameter needed to achieve these fits was usually ∼50%. Thus, despite the small effects observed in our signaling assay, the effects on the underlying biomolecular reactions are inferred to be substantial. We speculate that this might explain the evolutionary conservation of motifs that appear to be dispensable. This suggests that the pSLiMs are required for the efficiency of signaling, but they are not required for the reactions to occur. For mutants that showed large effects in our assay, the perturbed models tended to over predict the variation, which we believe is due to the simplicity of the model.

An example of a large effect mutation is the deletion of a known motif in Hot1(*30*). Consistent with a recent report that regulation of Hot1 by Casein Kinase (CK2) affects the stochasticity of the signaling response (*33*), we found that mutation of the known short motif in Hot1 also leads to a bimodal response, supporting the model that stochastic responses in this signaling pathway originate at the level of transcriptional regulation. To our knowledge, our results for the effects of mutations in a short motif in Hot1 is the first demonstration that short motifs in disordered regions can modulate the stochastic response of a signaling pathway (*20*, *36*).

Because we identified new motifs in the disordered regions of a well-characterized signaling pathway in yeast, our results suggest that most signaling pathways probably contain large numbers of unappreciated functional elements in disordered regions. We only investigated one type of osmotic stress at a single concentration and tested only single motif deletions, which overlook the possibility of cooperative interactions between weak motifs (*31*). Furthermore, HOG pathway components take part in other signaling networks. For example, the known SH3 binding site in Ste20 (*32*, *39*) was not identified, likely because this interaction is involved in mating signaling and not HOG signaling. Therefore, it’s likely that additional exploration will uncover functions for the pSLiMs that showed no effects in our assay.

If we can extrapolate from our results on the HOG pathway where we found 10 out of 29 (34%) pSLiMs had significant effects on pathway output. We predict at least 34% of pSLiMs found in other signaling pathways would also be functionally significant. As an example, we applied the same phyloHMM software to predict candidate motifs in another MAPK pathway in yeast and found 36 pSLiMs in the cell wall integrity pathway (supplementary table S1). We suggest that at least 12 (34%) of these are functional. An analysis of human proteins (*29*) identified 24 pSLiMs in the human canonical Wnt signaling pathway, of which we estimate 8 (34%) would be important for signaling. Taken together, this suggests that there is still a large number of short motifs to be discovered in well studied signaling pathways (*40*) and that comparative proteomics methods will be powerful tools to focus discovery efforts (*4*, *29*, *41*–*43*).

Here we have focused on short motifs because some of these can be identified using comparative proteomics approaches (*4*, *29*, *41*–*43*). However, we note that our comparative proteomics prediction of short conserved elements in the HOG pathway is likely to have missed important functional elements (*4*) - the number of unidentified motifs in disordered signaling molecules is likely even larger. Furthermore, In addition to short conserved SLiMs, there are many other types of functions that disordered regions may mediate in signaling pathways (*1*, *2*, *44*). Reliably identifying the currently uncharacterized functions of disordered regions will likely be critical to fully characterizing the mechanisms of signaling in human pathways. Indeed, approximately 22% of human disease mutations are now known to occur in IDRs (*45*), thereby implicating both IDRs and the undiscovered SLiMs (and other functional elements they contain) in any mechanistic understanding of human disease.

## Experimental Procedures

### Yeast strains and mutagenesis

HOG pathway flow cytometry assays were all performed in BY4741 (S288C-derivative strain: MATa his3Δ1 leu2Δ0 met15Δ0 ura3Δ0) lacking the Sho1-independent branch of the osmoresponse pathway (*ssk2*Δ*, ssk22*Δ) (*22*, *46*). The GFP-based reporter for the flow cytometry assay contained 800 bp of the native Stl1 promoter controlling expression of the yeast-enhanced yEGFP (*16*), integrated into the HO locus (*SSK2* Δ*::HisMX3 SSK22* Δ*::0; HO* Δ*::STL1pr-GFP,LEU2*). All subsequent deletions/mutations of the genomic sequence were performed using the *Delitto Perfetto* method for seamless mutagenesis of the yeast genome (*47*). The Positive control strain for the flow cytometry assay was created by deleting the conserved Sho1 Sh3 domain-binding site (Δ91-99) in Pbs2 (*26*). A secondary, more conservative control strain was created using double point mutations P96A and P99A (*26*) essential to the proline-rich Sh3 domain-binding site (PXXP).

### Pathway induction and flow cytometry assay

HOG pathway induction experiments were performed as follows: Replicate overnight cultures inoculated from single colonies were diluted 10-fold into the appropriate selective media and grown to mid-log phase (4 hours @ 30°C). Un-induced samples were taken at this point, and cells were pelleted and re-suspended in an equivalent volume of fixative (1X PBS, pH 7.0; 0.5% paraformaldehyde). Cultures were then induced by addition of NaCl to a final concentration of 0.4M and incubated @ 30°C for an additional 60 minutes. Induced samples were then taken and treated as above. All fixed samples were stored @ 4°C pending flow cytometry analysis (within 48 hours).

Reporter activation in fixed cells from all induction experiments was subsequently measured by flow cytometry. Flow cytometry analysis was performed on a BD FACSAria IIU High Speed Cell Sorter, incorporating 3 air-cooled lasers at 488, 633, and 407 nm wavelengths, and equipped with BD FACSDiva™ software (v. 6.1.3). The instrument was calibrated prior to each experiment using a blend of size-calibrated fluorescent beads. The GFP fluorescence was excited at 488 nm and collected through 530/30 nm bandpass filter. All samples were further diluted by a factor of 10, and a total of 50,000 events (cells) were counted for every sample within gated SSC and FSC populations that excluded doublets and small debris. Typical acquisition rates were 700-900 events/second. Initial flow cytometry data was analyzed using FLowJo software (v 10, OSX)

### Histogram data analysis

Data analysis of flow cytometry data was performed using in-house Matlab scripts. We used KL divergence to compare distributions. Because KL divergence is based on continuous probability distributions, to implement it on discrete data set, we used an approximation to estimate the KL divergence based on the empirical distribution (*48*).

### Comparison of mutant distributions to wild type distribution

We pooled WT data from the same experiment day (3 to 4 replicates) which we refer to as the WT pool. We then used this WT pool as the Q distribution to calculate KL divergence. Used in this way, the KL divergence can be thought of as the error one would make if the wild type distribution were used in place of a given mutant distribution. The KL divergences of each WT replicate from the WT pool of the same experiment day is calculated and grouped to estimate the variance in WT reporters. We then calculated the KL divergence of a mutant replicate from the WT pool of the same experiment day. The divergences of the same mutant from different experiment days are then grouped together. We tested if the mutants’ divergences from WT pools are different from WT’s divergences from WT pools with the Willcoxon Rank Sum test.

### A stochastic model of HOG transcriptional activation

We implemented the stochastic model (*21*) using SimBiology in MATLAB. The value of the stress parameter was set to 18, which minimizes the KL divergence from our wild-type pools under 0.4 M NaCl stress after 60 minutes. We modeled the effects of mutation by decreasing the value of 2 parameters, one that determines the activation of Hog1 by stress, the other determines the frequency that remodeler binds to DNA. We varied each parameter independently, in increments of 5% of the original value. In total, 40 perturbations (20 per parameter) were performed with 50,000 runs to represent 50,000 cells of each.

The distribution obtained for each perturbation was analyzed using the same method that we used to compare the mutant distributions to the wildtype distribution (described above) with the following modifications: first, a small lognormally distributed constant (mu, sigma = 2.5, 0.6) was added to the results of each run to represent basal gene expression (rather than normally distributed constant as assumed previously (*36*)). Second, the simulated data were linearly transformed such that two requirements are satisfied: 1) the maximum intensity was aligned to a replicate of WT data (which we subsequently excluded for comparisons of KL divergence). 2) The mean of unstressed simulated distribution was aligned to the mean of unstressed control for the particular experimental replicate being compared. The maximum and minimum intensity of the experimental replicate were introduced into the simulated distribution as a form of linear extension (*48*).

## Supplementary Figure Legend

**Figure S1.**
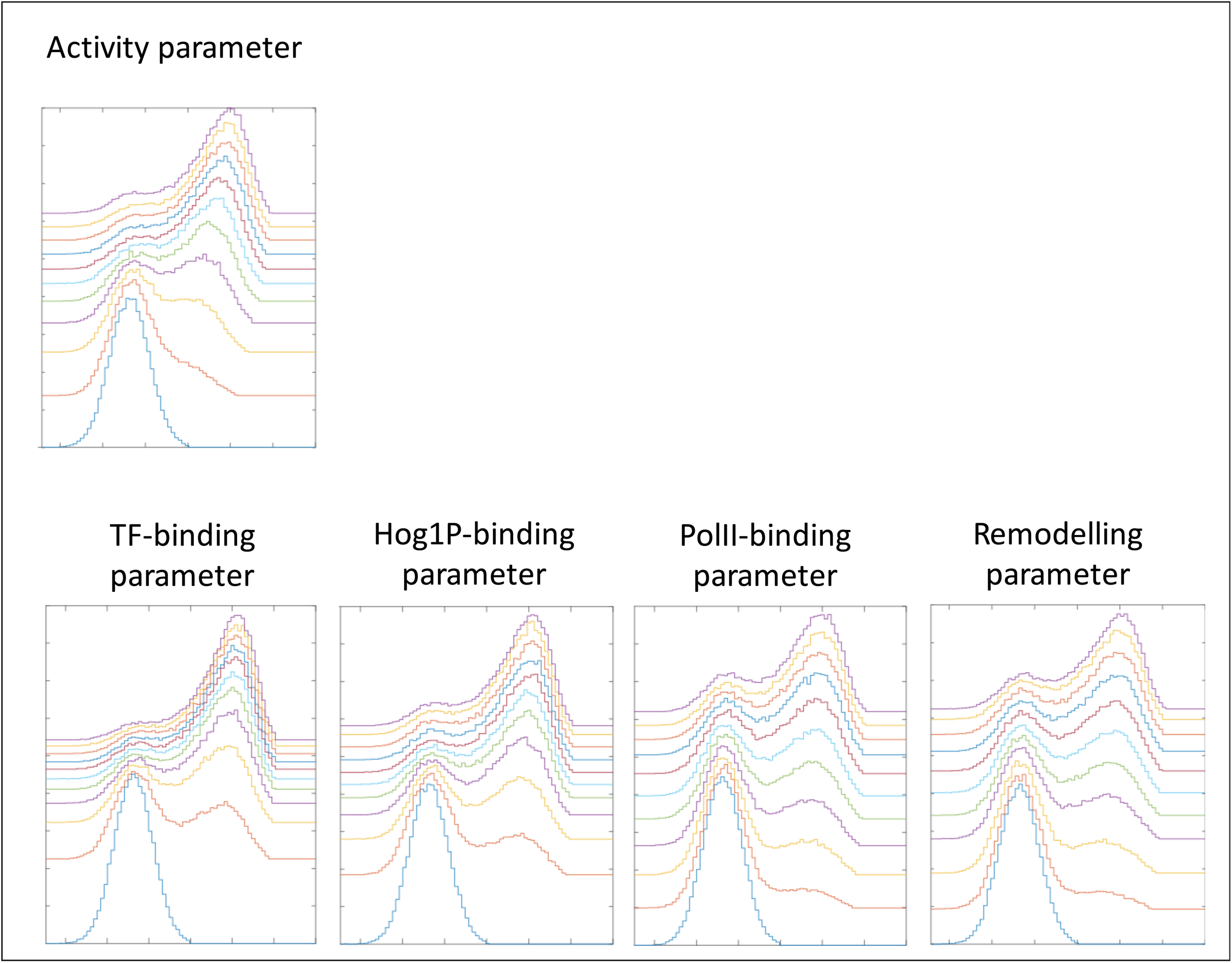
Effects of perturbations in model parameters fall into two groups. The activity parameter that affects the pathway activation steps upstream of the MAPK Hog1 (referred to as c_active in (*21*), top panel) affects both the proportion of cells that activate the pathway (right mode of distribution) as well as the maximum activity of the pathway (location of the right mode), such that as the parameter is reduced, fewer cells activate the pathway and those that do, do so to a lesser extent. The bottom panel shows that the four parameters that affect the slow transcriptional steps below Hog1 (referred to as c_tfon, c_hogon, c_remon, c_polon in (*21*)) affect the proportion of cells in the right mode, but not the location of the mode, so the distributions appear more bimodal. Each plot contains predictions of the model where the original parameter value has been permuted in the range from 0 to 50% of the original parameter value in increments of 5%.

## Supplementary Table Legends

**Table S1.**
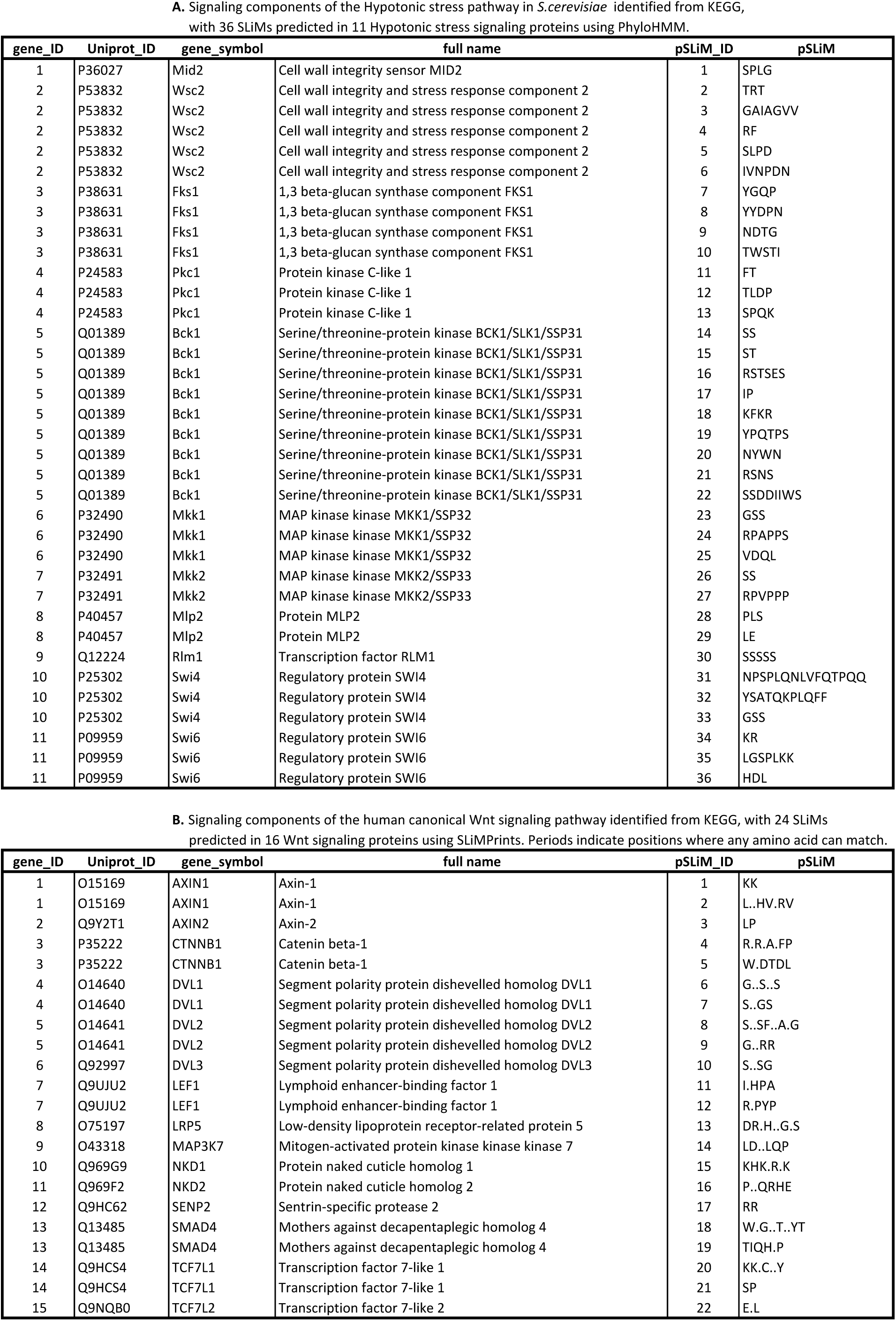
Predicted short linear motifs (pSLIMs) in the hypotonic stress and canonical Wnt signaling pathways. **A.** Signaling components of the hypotonic stress pathway in *S.cerevisiae* identified from KEGG, with 36 SLiMs predicted in 11 hypotonic stress signaling proteins using PhyloHMM. **B.** Signaling components of the human canonical Wnt signaling pathway identified from KEGG,with 24 SLiMs predicted in 16 Wnt signaling proteins using SLiMPrints. Periods indicate positions where any amino acid can match.

**Table S2.**
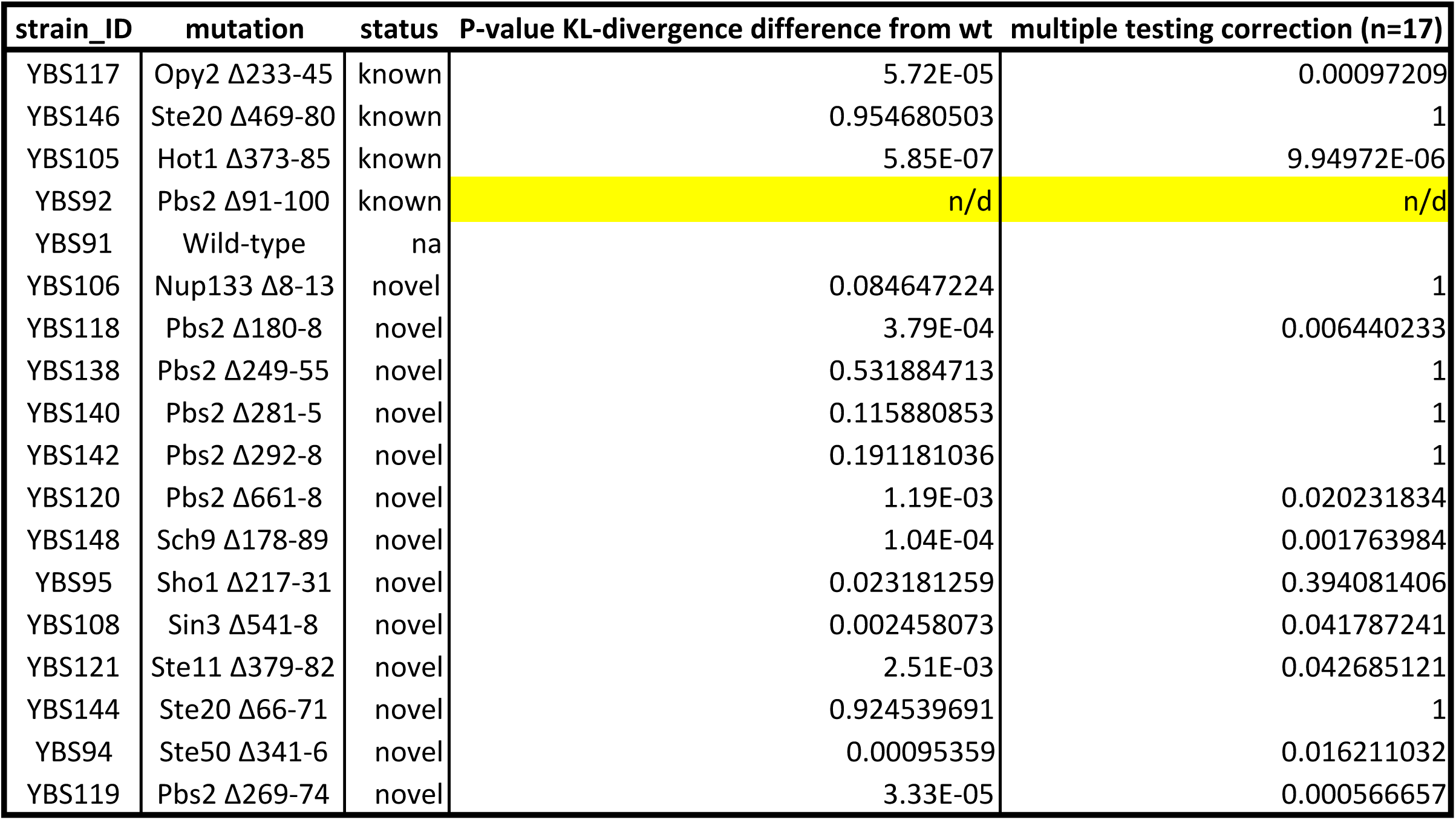
P-values from Willcoxon Rank Sum Tests comparing the distribution of KL divergence from mutants to wild-type pools.

## Acknowledgements

Thanks to Bianca Garcia and Dr. Alan Davidson for discussions and technical assistance. We thank Drs Audrey Gasch, Eric Weiss, Julie Forman-Kay, Yihan Lin and Muluye Liku for discussions and comments on the manuscript. BS and IH performed experiments. MLCM, BS and IH analyzed the data. All authors designed the experiments and wrote the manuscript. ANNB and TZ were funded by postgraduate scholarships from the Natural Sciences and Engineering Research Council of Canada (NSERC). This research was supported by a Canadian Institutes of Health research (grant MOP-119579) and by infrastructure grants from the Canadian Foundation for Innovation to AMM.

